# Detection of bifenthrin, bifenazate, etoxazole resistance in *Tetranychus urticae* collected from mint fields and hop yards using targeted sequencing and TaqMan approaches

**DOI:** 10.1101/2022.01.10.475553

**Authors:** Silas Shumate, Maggie Haylett, Brenda Nelson, Nicole Young, Kurt Lamour, Doug Walsh, Benjamin Bradford, Justin Clements

## Abstract

*Tetranychus urticae* (Koch) is an economically important pest of many agricultural commodities in the Pacific Northwest. Multiple miticides are currently registered for control including abamectin, bifenazate, bifenthrin, and extoxazole. However, populations of *Tetranychus urticae* have developed miticide resistance through multiple mechanisms, in many different growing regions. Producers of agricultural commodities where *Tetranychus urticae* infestations are problematic rely on integrated pest management tools to determine optimal control methods. Within this species multiple single nucleotide polymorphisms have been documented in different genes which are associated with miticide resistance phenotypes. The detection of these mutations through TaqMan qPCR has been suggested as a practical, quick, and reliable tool to inform agricultural producers of miticide resistance phenotypes present within their fields and have potential utility for making appropriate miticide application and integrated pest management decisions. Within this investigation we examined the use of a TaqMan qPCR-based approach to determine miticide resistance genotypes in field-collected populations of *Tetranychus urticae* from mint fields and hop yards in the Pacific Northwest of the United States and confirmed the results with a multiplex targeted sequencing. The results suggest the TaqMan approach accurately genotypes *Tetranychus urticae* populations collected from agricultural fields. The interpretation of the results, however, provide additional challenges for integrated pest management practitioners, including making miticide application recommendations where populations of *Tetranychus urticae* are a mix of resistant and wildtype individuals.

## 1. Introduction

*Tetranychus urticae* (Koch) is an economically relevant pest of multiple agricultural commodities in the Pacific Northwest, including hops and mint (1–2). *Tetranychus urticae* are small (1/50 inch long) mites that predominantly live and feed on the underside of plant leaves (3). This species is incredibly adaptable and can feed on more than 1,000 different plant species, including multiple cropping systems found in Idaho and Washington (1,4). The gnathosoma of *T. urticae* is complex, with chelicera that have been adapted to needle-like mouthparts for piercing and sucking of plant material, removing chlorophyll from plant cells, and causing extensive damage (5–7). The damaged caused by *T. urticae* can result in necrotic spots, yellowing, and loss of leaves that are fed upon (8–9). High infestations late in the growing season in the Pacific Northwest can result in economic loss of crops, including mint stands and hop yards. To control populations of *T. urticae* growers rely on both cultural integrated pest management practices along with miticide compounds (10).

To date numerous populations of *T. urticae* have developed resistance to over 96 different miticide chemistries, with over 140 publications documenting the development of resistance (11). Miticide resistance has been documented in Africa, Asia, Australia, Europe, North America, and South America (11). Currently, agricultural producers proactively monitor pest populations through arthropod scouting for pest population thresholds and apply foliar miticides when established thresholds are reached (12). If the miticide application is ineffective, growers can screen pest populations for miticide resistance using median lethal dose assays (LD_50_) or by sending pest populations to extension agencies or pest scouting agents to be screened. However, these approaches can take weeks to determine resistant phenotypes. Instead, the grower will usually make an informed decision and apply another pesticide treatment for control. Unfortunately, using the incorrect chemical or wrong concentration of the correct chemical may not provide adequate control and/or can further drive the development of miticide resistance.

Miticide resistance can develop through multiple mechanisms, including enhanced metabolic breakdown of miticides, target site insensitivity, and behavioral resistance (13–14). One of the most important mechanisms for miticide resistance development in *T. urticae* is target site insensitivity, where single nucleotide polymorphisms (SNPs) can result in an alteration of an amino acid sequence and the translated protein which, in-turn, binds the insecticide more weakly or not at all (13). These polymorphisms are non-synonymous and can spread through pest populations through natural selection when they result in beneficial traits (15). SNPs in multiple different genes have become associated with the development of miticides resistance to different miticide chemistries including bifenthrin, bifenazate, and etoxazole (16–19). Historically these mutations have been detected through DNA sequencing that can be analyzed for the presence of the polymorphism. However, multiple investigations have recently suggested using these polymorphisms as a diagnostic tool to inform agricultural growers of the miticide resistance status of individual field populations of *T. urticae*. This would allow the grower to make an informed decision into which miticide would adequately control their local *T. urticae* populations and may prevent further resistance development (20–21). Three of the most common polymorphisms studied are a mutation that results in the amino acid change of a glycine to a serine in the cytochrome b gene in the mitochondrial respiratory chain. This mutation has been shown to confer resistance to bifenazate, a carboxylic ester, in *T. urticae* (22). Another mutation at amino acid 1538 in the voltage gate channel protein s phenylalanine to isoleucine results in resistance to bifenthrin, a pyrethroid (16). Additionally, a mutation that changes isoleucine to phenylamine in the chitin synthase 1 gene has been shown to result in resistance to etoxazole, a narrow spectrum systemic acaricide (19). These mutations have been characterized by sequencing both susceptible and resistant populations of *T. urticae* and have provided in-depth information on the development of resistance to miticides in *T. urticae* (16–20, 22).

One potential approach for detecting resistance mutations is to use a TaqMan quantitative PCR genotyping assay (20–21). This approach allows for the detection of SNPs using two fluorescent probes; one labeled to detect the wildtype allele and the other to detect the allele with the mutation. The presence of the mutation can be quantified within a DNA sample and can be used to classify a population as wildtype (susceptible), heterozygous (susceptible/resistant), or resistant phenotype. This information can be used to determine the efficacy of a miticide before an application is ever applied. In the present study, we used a TaqMan qPCR approach to monitor resistant populations of *T. urticae* and examine the genotypes of *T. urticae* collected from mint fields and hop yards in the Pacific Northwest of the United States. We further confirmed our results with a multiplex PCR. Our results indicated that a TaqMan qPCR-based approach can quickly and accurately predict population allelic frequencies for polymorphisms that encode for resistance.

## 2. Materials and Methods

### 2.1 Data availability

All relevant data are contained within this paper and its supporting information files.

### 2.2 Ethics statement

This article does not contain studies with any human participants and no specific permits were required for collection or experimental treatment of *Tetranychus urticae* for the study described.

### 2.3 Tetranychus urticae Collections

*Tetranychus urticae* were collected from mint fields and hop yards in the Pacific Northwest during the summer of 2021. Briefly, *T. urticae* infested leaves were hand collected, placed in a sealed plastic bag, and transported in a cooler back to the Parma Research and Extension Center, Parma, Idaho or the Irrigated Agriculture Research and Extension Center (IAREC), Prosser, Washington. Samples were shared between research stations. A total of thirteen populations of *T. urticae* were collected from Washington and Idaho (9 populations from Washington and 4 populations from Idaho). Samples were placed in insect/mite proof mesh cages and housed on dry beans while DNA extraction and median lethal dose assays were conducted.

**Figure 1:**
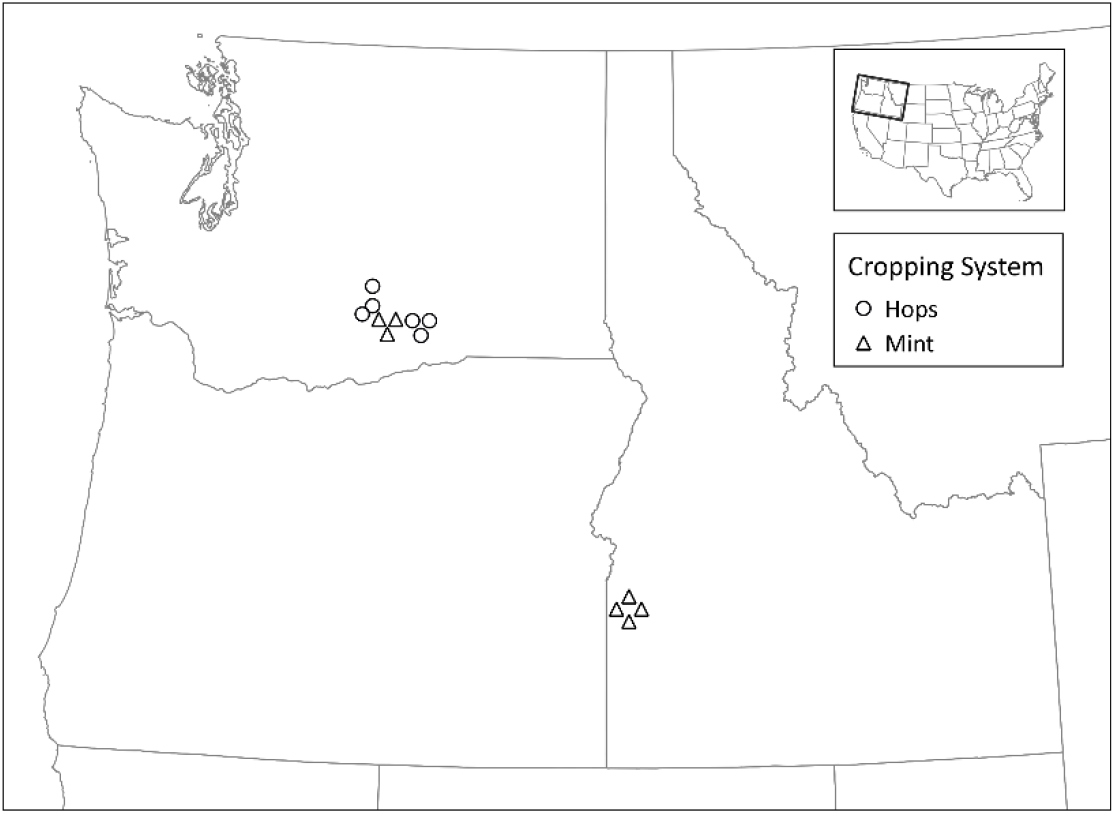
Field collection sites for *Tetranychus urticae* from mint fields and hop yards in the Pacific Northwest of the United States.

### 2.4 DNA isolation from *Tetranychus urticae*

*Tetranychus urticae* DNA was isolated using two different methods; the first was a Cethyl Trimethyl Ammonium Bromide (CTAB) method and the second was a Zymo Research Quick-DNA™ Tissue/ Insect Microperp kit. Heavily infested *T. urticae* leaves were removed from sealed cages and placed in a sealed petri dish to eliminate movement of mites. Mites were brushed from leaves with a mite brushing machine (Leedom Engineering, CA, USA) onto glasses plates. The mite brushing machine was cleaned before and after use and sterilized with UV light for 15 minutes. After samples of mites were collected, the glass plates were chilled at 4°C for 20 mins. The plate was observed under a dissecting microscope to confirm mite species and remove any possible contaminants. Mite samples were then scraped into 2 ml polypropylene tubes using two sterile razor blades. This method of removing the mites from the leaves was done for both extraction methods.

For the modified CTAB extraction protocol, aggregate samples of mites were placed in 2 ml DNAse/RNAse-free homogenate tubes (Biospec, OK, USA) with a single sterilized 6.4 mm and four 2.7 mm diameter glass bead (Biospec, OK, USA). Seven hundred and fifty μl of CTAB Extraction Buffer was added to each tube (OPS diagnostics, NJ, USA). Samples were homogenized for 2 minutes in a Mini-beadbetter-16 (Biospec, OK, USA). Tubes with homogenate were incubated at 60 °C in a water bath for 30 minutes. Following the incubation period, samples were centrifuged for 10 minutes at 14,000 x g and the supernatant was transferred to a new 1.5 ml tube. Five μl of RNase solution A (20 mg/ml, Fisher Scientific, MA, USA) was added and incubate at 37 °C for 20 minutes. Three hundred μl of chloroform/isoamyl alcohol (24:1) was added to each sample and vortexed for 5 seconds and samples were then centrifuged for 1 minute at 14,000 x g to separate the phases. The chloroform/isoamyl alcohol step was conducted twice. The upper aqueous phase was transferred to a new 1.5 ml microcentrifuge tube. DNA was precipitated by adding 500 μl cold isopropanol. Samples were left for 12 hours at −20 °C. Samples were then centrifuged at 14,000 x g for 10 minutes to pelletize DNA. Supernatant was decanted and discarded without disturbing the pellet, which was subsequently washed with 1 ml of ice cold 70% ethanol, and then vortexed and centrifuged at 14,000 x g for 10 minutes. Ethanol was decanted and excess ethanol was removed with a pipettor from the pellet. Samples were air dried in a sterile PCR cabinet for 15 min. DNA was dissolved in 100 μl RNase/DNase free H2O. DNA concentration was determined using a Nanodrop 2000. Samples were stored at −20 °C until multiplex and TaqMan processing. For the Zymo Research Quick-DNA™ Tissue/ Insect Microperp kit extraction method, aggregate samples of mites were added to ZR BashingBead™ Lysis Tube (2.0 mm) and 750 μl BashingBead™ buffer. The mites within the tubes were then homogenized for 6 minutes, after which Zymo Research Quick-DNA™ Tissue/ Insect Microprep kit methodology was followed, and DNA was eluted from the spin column with 20 μl of DNA Elution Buffer. *T. urticae* DNA concentration was determined using a Nanodrop 2000. Samples were stored at −20 °C until multiplex PCR and TaqMan qPCR processing.

### 2.5 Genotyping Quantitative PCR

TaqMan quantitative PCR was used to examine different allele frequencies within populations of *T. urticae* for mutations in G126S (Cytochrome b) (23), F1538I (Voltage gated sodium channel) (15), and I1017F (Chitin synthase 1) (18). Accession numbers for each reference gene can be found in **Supplemental File S1**. TaqMan based primers and probes were designed and purchased through Custom TaqMan® SNP Genotyping Assays Designs Tool (Thermo Fisher, MA, USA). Primer and probes can be found in **Supplemental File S1**. The TaqMan-based assay uses both FAM and VIC fluorescent probes, with the wildtype allele labeled with the VIC probe and the FAM probe for the mutation that is associated with the resistant allele. The TaqMan qPCR reaction was conducted in 25 μl reactions using the Applied Biosystems™ TaqMan™ Genotyping Master Mix (Applied Biosystems, MA, USA) and followed recommended procedures. Duplicate reactions were run at 95 °C for 10 min, followed by 95 °C for 15 s, and 60 °C for 60 s for a total of 40 cycles on a CFX 96 Bio-Rad machine (Applied Biosystems, MA, USA). Data was analyzed using Bio-Rad CFX Maestro Software (Applied Biosystems, MA, USA). Unknown calls were mapped to the closest phenotype. Negative template controls were run along with all samples as outline in Bio-Rad CFX software manual (Applied Biosystems, MA, USA).

### 2.6 Multiplex targeted-sequencing

Total DNA from each sample was sent to Floodlight Genomics, LLC. Floodlight Genomics used an optimized Hi-Plex approach to amplify and barcode targets in a single multiplex PCR reaction followed by sequencing on an Illumina HiSeqX device running a 2×150bp paired-end configuration, as previously described (24). Primers were designed to amplify five 150 bp regions to examine known SNPs previously correlated with miticide resistance in *T. urticae* (**Supplemental File S1**), G126S (Cytochrome b) (22), F1538I (Voltage gated sodium channel) (16), F1534S (Voltage gated sodium channel) (16), M918L (Voltage gated sodium channel) (16) and I1017F (Chitin synthase 1) (19). The sample-specific barcoded amplicons were sequenced on the Illumina HiSeq X platform according to the manufacturer’s directions. Floodlight Genomics delivered sample-specific raw DNA sequence reads as FASTQ files. Annotation of the raw reads was performed with Geneious Bioinformatics Software (Auckland, New Zealand). The reads were then mapped to reference sequences at 100% stringency to classify genotype.

### 2.7 Medium Lethal Dose Assay

Median lethal dose bioassays were conducted on 1 cm bean leaf disks with 4 replicate leaf disks per rate tested. Ten gravid adult female *T. urticae* were placed on each leaf disk. These mites were treated with acaricides at increasing rates. Bifenazate treatment rates were 0 (control), 3.12%, 6.25%, 12.5%, and 50% of the maximum label rates permitted of Vigilent 4SC (24 fluid oz per acre). Each bioassay arena for each of the rates tested was treated with 2 ml of the dilute solution. After 24 hours mites were evaluated for mortality. Mites were classified as dead when they failed to move more than their body length when prodded with a small paint brush. Moribund mites rarely recover and typically die within the next 24 hrs. The LC_50_ (lethal concentration that controls 50 % of the population) the 95% CI (confidence interval), the slope ± SEM (standard error of the mean), X^2^ and df were calculated with the software Polo Probit (LeOra Software LLC).

## 3. Results

### 3.1 TaqMan quantitative PCR detection of bifenazate (G126S), bifenthrin (F1538I), and etoxazole (I1017F) resistance

The TaqMan PCR genotyping assay detected mutations encoding for bifenazate (G126S), bifenthrin (F1538I), and etoxazole (I1017F) resistance in samples of *T. urticae* collected in mint and hops in Washington and Idaho (**Table 1**). Within the populations that were examined, we observed that no population had a completely resistant genotype that was fixed for bifenazate (G126S) resistance. Most populations collected from mint were identified as wildtype susceptible, while most populations from hops were heterozygous and composed of individuals with both wildtype and resistant alleles. For bifenthrin (F1538I), we observed most populations of mites collected were heterozygous. One population collected in mint from Idaho identified as wildtype susceptible and another population collected from mint in Idaho had a resistant genotype that was fixed for bifenthrin (F1538I). Etoxazole (I1017F) resistance was split regionally, with populations of mites collected in Idaho expressing wildtype phenotypes and populations of mites collected in Washington expressing heterozygous or resistance phenotypes.

**Table 1.** TaqMan assay of *T. urticae* collected from hops and mint in the Pacific Northwest.

| Pop. | Cropping System | Growing Region | G126S<br>(Bifenazate) | F1538I<br>(Bifenthrin) | I1017F<br>(Etoxazole) |
| --- | --- | --- | --- | --- | --- |
|  |  |  | TaqMan | TaqMan | TaqMan |
| 1 | Hops | Washington | Heterozygous | Heterozygous | Resistant |
| 2 | Hops | Washington | Heterozygous | Heterozygous | Resistant |
| 3 | Hops | Washington | Wildtype | Heterozygous | Heterozygous |
| 4 | Hops | Washington | Heterozygous | Heterozygous | Heterozygous |
| 5 | Hops | Washington | Heterozygous | Heterozygous | Heterozygous |
| 6 | Hops | Washington | Heterozygous | Heterozygous | Heterozygous |
| 7 | Mint | Idaho | Wildtype | Heterozygous | Wildtype |
| 8 | Mint | Idaho | Wildtype | Heterozygous | Wildtype |
| 9 | Mint | Idaho | Wildtype | Resistant | Wildtype |
| 10 | Mint | Idaho | Heterozygous | Wildtype | Wildtype |
| 11 | Mint | Washington | Wildtype | Heterozygous | Heterozygous |
| 12 | Mint | Washington | Wildtype | Heterozygous | Heterozygous |
| 13 | Mint | Washington | Wildtype | Heterozygous | Heterozygous |

### 3.2 Targeted-sequencing and detection of bifenthrin (F1538I, F1535I, M918L) and etoxazole (I1017F) resistance

A multiplex targeted-sequencing approach provided additional insight into the relative frequency of the SNPs within each population. The multiplex assay provides read counts that can be correlated to the mutation within a population. Two additional mutations from the TaqMan qPCR were examined within this analysis; F1534S (Bifenthrin) and M918L (Bifenthrin). The mutation that encoded for bifenazate (G126S) was not successfully amplified in any of the samples. Within the analysis of the targeted-sequencing data, we examined two additional SNPs which encoded for bifenthrin resistance, but neither mutation was found within any of the populations examined. Further, this technology allowed us to examine the relative fixation of genotypes within a sample population. In population 1 (collected from hops in Washington), we noted that 98.2% of the reads encoded for resistance to etoxazole (I1017F), suggesting a highly fixed mutation within this population, while for population 9 (collected from mint in Idaho) we noted that 91.7% of the population had resistance to bifenthrin (F1538I). This suggests that 8.3% of the reads were still wildtype and the population has not become homozygous for the resistant alleles. This information provides additional insight into the relative frequency of the resistant phenotype within field collected populations which may be useful in choosing effective miticides and integrated pest management approaches.

The results from the TaqMan qPCR and targeted-sequencing assays were compared for F1538I (Bifenthrin) and I1017F (Etoxazole) to confirm results (**Table 3**). The results from both methods had high agreement. The targeted-sequencing approach was more sensitive and provided accurate data on the proportion of the presence of the frequency of the allele. However, the TaqMan assay was able to provide accurate calls in 23 out of 24 of the runs. The TaqMan assay predicted population 2 (collected from hops in Washington) as a completely resistant population when, in fact, there was a high proportion of wildtype alleles (17.45%). As such, the population should be considered heterozygous.

**Table 2.**
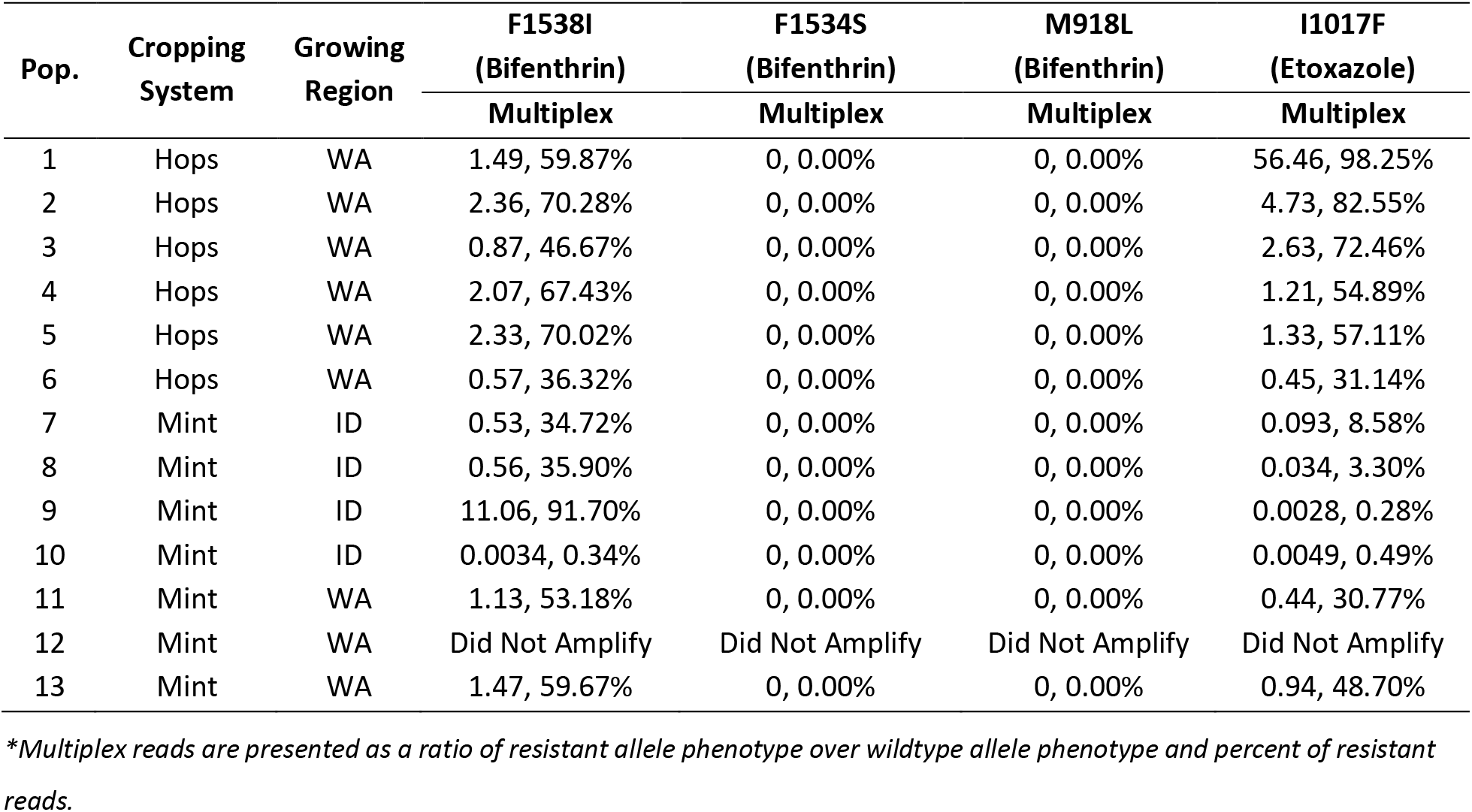
Multiplex PCR assay of *T. urticae* collected from hops and mint in the Pacific Northwest.

| Pop. | Cropping System | Growing Region | F1538I<br>(Bifenthrin) | F1534S<br>(Bifenthrin) | M918L<br>(Bifenthrin) | I1017F<br>(Etoxazole) |
| --- | --- | --- | --- | --- | --- | --- |
|  |  |  | Multiplex | Multiplex | Multiplex | Multiplex |
| 1 | Hops | WA | 1.49, 59.87% | 0, 0.00% | 0, 0.00% | 56.46, 98.25% |
| 2 | Hops | WA | 2.36, 70.28% | 0, 0.00% | 0, 0.00% | 4.73, 82.55% |
| 3 | Hops | WA | 0.87, 46.67% | 0, 0.00% | 0, 0.00% | 2.63, 72.46% |
| 4 | Hops | WA | 2.07, 67.43% | 0, 0.00% | 0, 0.00% | 1.21, 54.89% |
| 5 | Hops | WA | 2.33, 70.02% | 0, 0.00% | 0, 0.00% | 1.33, 57.11% |
| 6 | Hops | WA | 0.57, 36.32% | 0, 0.00% | 0, 0.00% | 0.45, 31.14% |
| 7 | Mint | ID | 0.53, 34.72% | 0, 0.00% | 0, 0.00% | 0.093, 8.58% |
| 8 | Mint | ID | 0.56, 35.90% | 0, 0.00% | 0, 0.00% | 0.034, 3.30% |
| 9 | Mint | ID | 11.06, 91.70% | 0, 0.00% | 0, 0.00% | 0.0028, 0.28% |
| 10 | Mint | ID | 0.0034, 0.34% | 0, 0.00% | 0, 0.00% | 0.0049, 0.49% |
| 11 | Mint | WA | 1.13, 53.18% | 0, 0.00% | 0, 0.00% | 0.44, 30.77% |
| 12 | Mint | WA | Did Not Amplify | Did Not Amplify | Did Not Amplify | Did Not Amplify |
| 13 | Mint | WA | 1.47, 59.67% | 0, 0.00% | 0, 0.00% | 0.94, 48.70% |

\*Multiplex reads are presented as a ratio of resistant allele phenotype over wildtype allele phenotype and percent of resistant reads.

**Table 3.** Comparison between TaqMan qPCR and multiplex PCR results in *T. urticae* collected from hops and mint in the Pacific Northwest.

| Pop. | Cropping System | Growing Region | F1538I (Bifenthrin) |  | I1017F (Etoxazole) |  |
| --- | --- | --- | --- | --- | --- | --- |
|  |  |  | TaqMan | Multiplex | TaqMan | Multiplex |
| 1 | Hops | Washington | Heterozygous | 1.49, 59.87% | Resistant | 56.46, 98.25% |
| 2 | Hops | Washington | Heterozygous | 2.36, 70.28% | Resistant | 4.73, 82.55% |
| 3 | Hops | Washington | Heterozygous | 0.87, 46.67% | Heterozygous | 2.63, 72.46% |
| 4 | Hops | Washington | Heterozygous | 2.07, 67.43% | Heterozygous | 1.21, 54.89% |
| 5 | Hops | Washington | Heterozygous | 2.33, 70.02% | Heterozygous | 1.33, 57.11% |
| 6 | Hops | Washington | Heterozygous | 0.57, 36.32% | Heterozygous | 0.45, 31.14% |
| 7 | Mint | Idaho | Heterozygous | 0.53, 34.72% | Wildtype | 0.093, 8.58% |
| 8 | Mint | Idaho | Heterozygous | 0.56, 35.90% | Wildtype | 0.034, 3.30% |
| 9 | Mint | Idaho | Resistant | 11.06, 91.70% | Wildtype | 0.0028, 0.28% |
| 10 | Mint | Idaho | Wildtype | 0.0034, 0.34% | Wildtype | 0.0049, 0.49% |
| 11 | Mint | Washington | Heterozygous | 1.13, 53.18% | Heterozygous | 0.44, 30.77% |
| 12 | Mint | Washington | Heterozygous | Did Not Amplify | Heterozygous | Did Not Amplify |
| 13 | Mint | Washington | Heterozygous | 1.47, 59.67% | Heterozygous | 0.94, 48.70% |

\*Multiplex reads are presented as a ratio of resistant allele phenotype over wildtype allele phenotype and percent of resistant reads.

### 3.3 Median Lethal Dose Assay

Median lethal dose assays were conducted on 6 populations of mites: 3 collected from hops and 3 collected from mint in Washington (**Table 4**). The populations of mites collected from hops were all heterozygous for bifenazate resistance, while the populations collected from mint all had susceptible phenotypes. The results from the LD_50_ experiment on field collected mites from hops and mint suggested that none of the examined populations had a resistant phenotype compared to the laboratory population. The results from the LD_50_ assay suggest that even though the populations from hops had heterozygous resistant phenotypes, the expression of that allele does not accurately predict whether the population has a resistant phenotype. The populations collected in mint had a more resistant phenotype than the mites collected in hops when exposed to bifenazate. The results suggest that using this specific marker alone is not a significant predicator of resistance to bifenazate.

**Table 4.** Median lethal dose assay to bifenazate from *T. urticae* collected in mint fields and hop yards from Washington.

| Population | LD<sub>50</sub> (ppm) | 95% CI | Slope ± SEM | χ<sup>2</sup> | df | RR* |
| --- | --- | --- | --- | --- | --- | --- |
| Laboratory susceptible | 0.53 | 0.46-0.66 | 3.35±0.53 | 7.48 | 14 | 1 |
| 1 - Hops | 1.23 | 0.42-2.01 | 1.19±0.27 | 14.92 | 14 | 2.22 |
| 2 - Hops | 0.9 | 0.17-1.63 | 1.07±0.28 | 14.39 | 14 | 1.64 |
| 5 - Hops | 2.17 | 1.35-3.16 | 1.53±0.28 | 15.84 | 14 | 3.94 |
| Laboratory susceptible | 1.3 | 0.47-2.12 | 1.34±0.28 | 27.3 | 14 | 1 |
| 11 - Mint | 2.13 | 1.74-2.60 | 3.64±0.56 | 15.6 | 14 | 1.64 |
| 12 - Mint | 9.51 | 4.68-92.48 | 0.91±0.23 | 21.1 | 14 | 4.46 |
| 13 - Mint | 8.66 | 5.09-90.77 | 0.76±0.23 | 12.9 | 14 | 4.06 |

## 4. Discussion

In this investigation we examined five mutations in field populations of twospotted spider mites using two molecular diagnostic approaches. These mutations have been previously linked to miticide resistance phenotypes and include the mutation G126S in cytochrome b, F1538I, F1534S, M918L, in a voltage gated sodium channel, and I1017F in the chitin synthase 1 gene, and have all been shown to confer varying levels of resistance to commonly used miticides. We examined these markers using two different diagnostic approaches: a TaqMan qPCR and targeted-sequencing. The TaqMan qPCR has been suggested as a molecular tool for growers to monitor resistance status of *T. urticae* (20–21) and we set out to determine if a TaqMan-based approach would be a quick and reliable tool for growers to provide information regarding resistant phenotypes of local populations. Our findings confirm its utility to detect mutations in twospotted spider mites and further demonstrates that it can be used for field-collected populations. We also confirmed the TaqMan results were accurate using the complementary (and, in some cases, more sensitive) targeted-sequencing approach. The targeted-sequencing provided insight into the relative proportion of allele frequencies that encodes for resistance within a population of *T. urticae* and allowed us to test if the TaqMan qPCR-based approach provides similar results. The findings of this investigation suggest that a TaqMan based approach can provide quick and accurate data from field collected population of *T. urticae*. However, how a grower would interpret the results in the context of pest management decisions needs to be further addressed.

Using established primers and probes, a molecular laboratory can quickly assess the presence of SNPs that are associated with miticide resistance using a TaqMan qPCR approach. This method includes sampling *T. urticae*, removing mite samples from leaves, extracting DNA, and running the TaqMan assay, which can all be done within roughly 4 hours. However, understanding the results can be more challenging, including determining appropriate thresholds and actions for integrated pest management approaches. Determining resistance for a heterozygous population, where a portion of the population has the resistant allele and others do not, may prove challenging using the TaqMan assay alone. More data regarding the frequency of known resistance mutations may provide a more comprehensive picture of the resistance status of the population of *T. urticae*. For this, we examined the use of Illumina-based targeted-sequencing, which provided a reasonable estimate of the overall frequency of the mutation within a population. Using the same primers to detect both the wildtype and resistant alleles, we can deduce that the amplification frequency will be the same for each allele and the total number of reads for each allele can be used to estimate the frequency within a population. Both approaches are subject to error if there are unknown mutations in primer binding sites. Nonetheless, assessing the allele frequencies within a population sample may prove valuable to assist a grower in determining whether to apply a particular miticide. While the targeted-sequencing is more sensitive for assessing allele frequencies, it takes a significantly longer time to generate the data (amplifications, Illumina library prep and sequencing), which can be problematic for growers. The TaqMan qPCR and targeted-sequencing, however, provided similar results, with 23 of the 24 of the runs in agreement between the two methods.

Genotypic data generated from the TaqMan qPCR and targeted-sequencing provides significant insight on resistance phenotypes of *T. urticae* populations and miticide input within Idaho and Washington. A pyrethroid (bifenthrin) has never been registered for use on mint in the Pacific Northwest. The TaqMan qPCR and targeted-sequencing data for F1538I (Bifenthrin) suggested that most mint field populations of mites have some proportion of a resistance phenotype for Bifenthrin. Bifenthrin is used commonly in silage corn (25) which is grown intensely around mint fields in Idaho and Washington. In the fall, corn fields are treated routinely with pyrethroids in during harvest and *T. urticae* could then migrate into the adjacent mint fields. This movement between fields could explain why we see the markers that encode for resistance to bifenthrin in mites collected in mint, even when the compound is not currently used in mint fields. The data for G126S (Bifenazate) suggests that most populations of *T. urticae* from hops have a heterozygous phenotype, while mutations representing a resistant phenotype were rarely observed in *T. urticae* collected in mint in Idaho or Washington. Bifenazate is applied rarely to mint but heavily used in hops in the Pacific Northwest, which may explain the frequency of the resistant alleles found in this investigation. Etoxazole has cross-resistance with both hexythiozox and clofentazine. Washington mint growers have applied hexythiozox routinely for the past decade and the hop growers are applying both hexythiozox and etoxazole, yearly, in the hop yards in Washington. The phenotypic data for etoxazole matched prior expectations in Idaho and Washington, where we observed resistant phenotypes in hops yards in Washington and no resistant phenotype in mint fields in Idaho where etoxazole is not commonly used.

To confirm the results of the genotypic assays, we performed an LD_50_ assessment on bifenazate resistance. We decided to confirm resistance to one of the chemicals we explored, bifenazate, which can be predicted with the mutation G126S in the cytochrome b gene. The genotypic data suggested that *T. urticae* collected from hop yards and mint fields from Washington would demonstrate different resistant phenotypes. The LD_50_ assessment for *T. urticae* collected from mint fields demonstrated a higher LD_50_ value than *T. urticae* collected from hop yards. The results of the LD_50_ study suggested the exact opposite of the genetic data, with populations collected from mint fields demonstrating slightly higher bifenazate resistance than hop yards. Recent studies agree with our findings and suggest that the G126S mutation may not be a good indicator that a population is resistant to bifenazate (23). It is likely that multiple markers will be required to accurately detect resistance in field populations and further that the G126S mutation is not an optimal indicator of resistance status.

In this investigation we examined the use of using two different molecular approaches to determine the resistance genotypes present in *T. urticae* collected from mites and hops in the Pacific Northwest and infer their resistance phenotypes. The data suggests that TaqMan qPCR can be used to quickly genotype *T. urticae* collected from the field. However, the interpretation of the data might pose problems and concerns for integrated pest management decisions, including developing appropriate resistance allele frequency cutoffs for miticide applications with populations that have a heterozygous genotype. We further demonstrated that the presence the G126S mutation may not accurately predict bifenazate resistance and should not be used to predict actual resistance profiles. While these genetic approaches are constantly evolving based on our knowledge of putative and nearby polymorphisms (crucial for a stable assay), further work is needed to validate the importance of specific mutations and the metrics used for determining overall resistance profiles within a population to provide growers an accurate estimate of miticide resistance before it can be established as a useful pest management tool.

## 5. Acknowledgements

This research was supported in part by funding from United States Department of Agriculture Specialty Crop Initiative 2021-51181-35901, awarded to DW and JC, and a Mint Research Council grant awarded to DW and JC.

## 7. Supporting information

**Supplemental File S1: TaqMan primers and probes**

## References

1. Wu M, Adesanya AW, Morales MA, Walsh DB, Lavine LC, Lavine MD, Zhu F. Multiple acaricide resistance and underlying mechanisms in Tetranychus urticae on hops. Journal of Pest Science. 2019 Mar;92(2):543–55.

2. Adesanya AW, Franco E, Walsh DB, Lavine M, Lavine L, Zhu F. Phenotypic and genotypic plasticity of acaricide resistance in populations of Tetranychus urticae (Acari: Tetranychidae) on peppermint and silage corn in the Pacific Northwest. Journal of economic entomology. 2018 Dec 14;111(6):2831–43.

3. Rioja C, Zhurov V, Bruinsma K, Grbic M, Grbic V. Plant-herbivore interactions: a case of an extreme generalist, the two-spotted spider mite Tetranychus urticae. Molecular Plant-Microbe Interactions. 2017 Dec 10;30(12):935–45.

4. Agrawal AA. Host-range evolution: adaptation and trade-offs in fitness of mites on alternative hosts. Ecology. 2000 Feb;81(2):500–8.

5. Fasulo TR, Denmark HA. Twospotted Spider Mite, Tetranychus urticae Koch (Arachnida: Acari: Tetranychidae). EDIS. 2003;2003(15).

6. Bensoussan N, Santamaria ME, Zhurov V, Diaz I, Grbić M, Grbić V. Plant-herbivore interaction: dissection of the cellular pattern of Tetranychus urticae feeding on the host plant. Frontiers in Plant Science. 2016 Jul 27;7:1105.

7. Park YL, Lee JH. Leaf cell and tissue damage of cucumber caused by twospotted spider mite (Acari: Tetranychidae). Journal of Economic Entomology. 2002 Oct 1;95(5):952–7.

8. Weihrauch F. Evaluation of a damage threshold for two-spotted spider mites, Tetranychus urticae Koch (Acari: Tetranychidae), in hop culture. Annals of applied biology. 2005 Jul;146(4):501–9.

9. Fasulo TR, Denmark HA. Twospotted Spider Mite, Tetranychus urticae Koch (Arachnida: Acari: Tetranychidae). EDIS. 2003;2003(15).

10. UC IPM. Two spotted Spider Mites. Accessed December 12th, 2021. Available at https://www2.ipm.ucanr.edu/agriculture/floriculture-and-ornamental-nurseries/Twospotted-spider-mite/

11. Michigan State University. Arthropod Pesticide Resistance Database. Accessed December 12th, 2021. Available at https://www.pesticideresistance.org/display.php?page=species&arId=536

12. Foster R, Flood BR. Vegetable insect management. 2005.

13. Van Leeuwen T, Vontas J, Tsagkarakou A, Tirry L. Mechanisms of acaricide resistance in the two-spotted spider mite Tetranychus urticae. InBiorational control of arthropod pests 2009 (pp. 347–393). Springer, Dordrecht.

14. Penman DR, Chapman RB, Bowie MH. Selection for behavioral resistance in twospotted spider mite (Acari: Tetranychidae) to flucythrinate. Journal of economic entomology. 1988 Feb 1;81(1):40–4.

15. Yang Z, Nielsen R. Estimating synonymous and nonsynonymous substitution rates under realistic evolutionary models. Molecular biology and evolution. 2000 Jan 1;17(1):32–43.

16. Nyoni BN, Gorman K, Mzilahowa T, Williamson MS, Navajas M, Field LM, Bass C. Pyrethroid resistance in the tomato red spider mite, Tetranychus evansi, is associated with mutation of the para-type sodium channel. Pest management science. 2011 Aug;67(8):891–7.

17. Demaeght P, Osborne EJ, Odman-Naresh J, Grbić M, Nauen R, Merzendorfer H, Clark RM, Van Leeuwen T. High resolution genetic mapping uncovers chitin synthase-1 as the target-site of the structurally diverse mite growth inhibitors clofentezine, hexythiazox and etoxazole in Tetranychus urticae. Insect biochemistry and molecular biology. 2014 Aug 1;51:52–61.

18. Kwon DH, Kang TJ, Kim YH, Lee SH. Phenotypic-and genotypic-resistance detection for adaptive resistance management in Tetranychus urticae Koch. PloS one. 2015 Nov 6;10(11):e0139934.

19. Van Leeuwen T, Demaeght P, Osborne EJ, Dermauw W, Gohlke S, Nauen R, Grbić M, Tirry L, Merzendorfer H, Clark RM. Population bulk segregant mapping uncovers resistance mutations and the mode of action of a chitin synthesis inhibitor in arthropods. Proceedings of the National Academy of Sciences. 2012 Mar 20;109(12):4407–12.

20. Ilias A, Vassiliou VA, Vontas J, Tsagkarakou A. Molecular diagnostics for detecting pyrethroid and abamectin resistance mutations in Tetranychus urticae. Pesticide biochemistry and physiology. 2017 Jan 1;135:9–14.

21. Mavridis K, Papapostolou KM, Riga M, Ilias A, Michaelidou K, Bass C, Van Leeuwen T, Tsagkarakou A, Vontas J. Multiple TaqMan qPCR and droplet digital PCR (ddPCR) diagnostics for pesticide resistance monitoring and management, in the major agricultural pest Tetranychus urticae. Pest Management Science. 2021 Sep 3.

22. Van Leeuwen T, Vanholme B, Van Pottelberge S, Van Nieuwenhuyse P, Nauen R, Tirry L, Denholm I. Mitochondrial heteroplasmy and the evolution of insecticide resistance: non-Mendelian inheritance in action. Proceedings of the National Academy of Sciences. 2008 Apr 22;105(16):5980–5.

23. Xue W, Wybouw N, Van Leeuwen T. The G126S substitution in mitochondrially encoded cytochrome b does not confer bifenazate resistance in the spider mite Tetranychus urticae. Experimental and Applied Acarology. 2021 Dec;85(2):161–72.

24. Nguyen-Dumont T, Pope BJ, Hammet F, Southey MC, Park DJ. A high-plex PCR approach for massively parallel sequencing. Biotechniques. 2013 Aug;55(2):69–74.

25. Ostile K, Managing Two-Spotted Spider Mites on Corn. University of Minnesota Extension.

