## Supplemental File for "Detection of bifenthrin, bifenazate, etoxazole resistance in *Tetranychus urticae* collected from mint fields and hop yards using targeted sequencing and TaqMan approaches"

Supplemental Table 1: Primers and probes for the TaqMan genotyping assay and primers for the multiplex PCR assay

| Assay | Mutation | Forward Primer Seq. | Reverse Primer Seq. | Reporter 1 Sequence | Reporter 2 Sequence | Accession # |
|---|---|---|---|---|---|---|
| TaqMan | F1538I | ACAACCAGTTTATGAAAATAGTATTCTGATGTACTT | CACCTCCTTTCTTTTTTTGTTCATTAAAATTATCAATAATG | ATTTTTGGCTCTTTTTTCACAC | TTTTGGCTCTTTTATCACAC | JN881331.1 |
| TaqMan | G126S | AGGATCCGCTTTTATTGGGT | AGTAATAACTGTTGCTCCCCAAAAAGAT | ATGTTTTACCTTGAGGACAAA | TGTTTTACCTTGAAGACAAA | EU556749.1 |
| TaqMan | I1017F | GCTTCATCCACAAGAGTTTCACTGT | ACGTTCAAGTTGACCAGAGAATAGAT | ATTTCCTTTCGATTCCATG | ATTTCCTTTCGTTTCCATG | tetur03g08510 |
| Multiplex | M918L | GAAGGTGTTCGAGGTCTTTCA | GATGGCCAGGACAAAGGTTA | NA | NA | JN881331.1 |
| Multiplex | F1534S | TGGACAATTATTATGGACCATGCAA | GACTCCAATAAACAGGTTAAGTGTGAAA | NA | NA | JN881331.1 |
| Multiplex | F1538I | AACAACCAGTTTATGAAAATAGTATTCTGATGTACTTA | CACCTCCTTTCTTTTTTTGTTCATTAAAATTATCAATAATG | NA | NA | JN881331.1 |
| Multiplex | G126S | AGGATCCGCTTTTATTGGGTATG | TCAACGGAAAATCTTCCCCAAAC | NA | NA | EU556749.1 |
| Multiplex | I1017F | TGCTTCATCCACAAGAGTTTCA | ACGTTCAAGTTGACCAGAGAATAG | NA | NA | tetur03g08510 |
